# Motor nuclei innervating eye muscles spared in mouse model of SOD1-linked ALS

**DOI:** 10.1101/304857

**Authors:** Eleanor V. Thomas, Maria Nagy, Hongyu Zhao, Wayne A. Fenton, Arthur L. Horwich

## Abstract

The eye muscles of humans with either inherited or sporadic forms of ALS are relatively unaffected during disease progression, a function of sparing of the cranial nerve motor neurons supplying them. Here we observe that cranial nerve nuclei are also spared in a mouse model of inherited SOD1-linked ALS. We examined the cranial nerve motor nuclei in a mouse strain, G85R SOD1YFP, which carries a high copy transgene encoding a mutant human SOD1-YFP fusion protein, that exhibits florid YFP-fluorescent aggregates in spinal cord motor neurons and paralyzes by 6 months of age. We observed in the cranial nerve nuclei that innervate the eye, 3N (oculomotor), 4N (trochlear), and 6N (abducens), that there was little (4N, 6N) or no (3N) aggregation, in comparison with other motor nuclei, 5N (trigeminal), 7N (facial), and 12N (hypoglossal), in the latter two of which florid aggregation was observed. Correspondingly, the number of ChAT positive motor neurons in 3N of G85R SOD1YFP relative to that in 3N of ChAT-EGFP mice showed that there was no loss of motor neurons over time, whereas in 12N there was progressive loss of motor neurons, amounting to a loss of ∼30% from G85R SOD1YFP by end-stage. Thus, the sparing of extraocular motor neurons as occurs in humans with ALS appears to be replicated in our SOD1-linked ALS mouse strain, supporting the validity of the mouse model for studying this aspect of selective motor system loss in ALS. Comparisons of extraocular motor neurons (e.g., from 3N), resistant to ALS pathology, with other cranial motor neurons (e.g., from 12N), sensitive to such pathology, may thus be of value in understanding mechanisms of protection vs. susceptibility of motor neurons.

---

The sparing of eye movements in ALS patients is well-known and has been related to lack of involvement of the motor nuclei innervating the extraocular muscles, namely, cranial nerve nuclei 3N (oculomotor), 4N (trochlear), and 6N (abducens) (ref. 1 and references therein). We asked whether our mouse model of ALS, bearing a high copy number transgene of a human genomic clone of G85R mutant SOD1 (including the human promoter) fused in its last exon with YFP (2,3), might likewise exhibit sparing of the same cranial nerve motor nuclei as compared with the other cranial motor nuclei. Notably, spinal cord motor neurons in our mouse strain exhibit large YFP fluorescent aggregates by 1 month of age, which increase in number subsequently and are associated with motor neuron cell death, amounting to 50% loss by 3 months of age and 75% loss by end-stage (3). The mice exhibit lower extremity symptoms by 3 months of age - twitching or pulling in when picked up by the tail – and uniformly paralyze by 6 months of age (4). We thus examined the cranial nerve nuclei at these two times, in each case examining 3 mice. At both 3 months and 6 months (end-stage), we observed no aggregation in ChAT positive motor neurons in 3N (see representative images in Fig.1A and 1B, and quantitation in bar graphs below and Table 1). In particular, 3N exhibited no aggregates in greater than 500 ChAT-positive motor neurons examined, ∼200 examined from three mice at 3 months and ∼300 examined from three mice at end-stage. The two other extraocular nuclei, 4N and 6N, exhibited rare aggregates at 3 months of age and essentially none at end-stage (see Fig. 1 bar graphs and Table 1). By contrast, 7N and 12N exhibited florid aggregation at both times (Fig. 1A, 1B, and Table 1), affecting ∼20-30% of neurons, whereas 5N was less affected, exhibiting a level of aggregation resembling that of 4N and 6N at 3 months, but greater than 3N, 4N, and 6N at end-stage.

**Fig.1.**
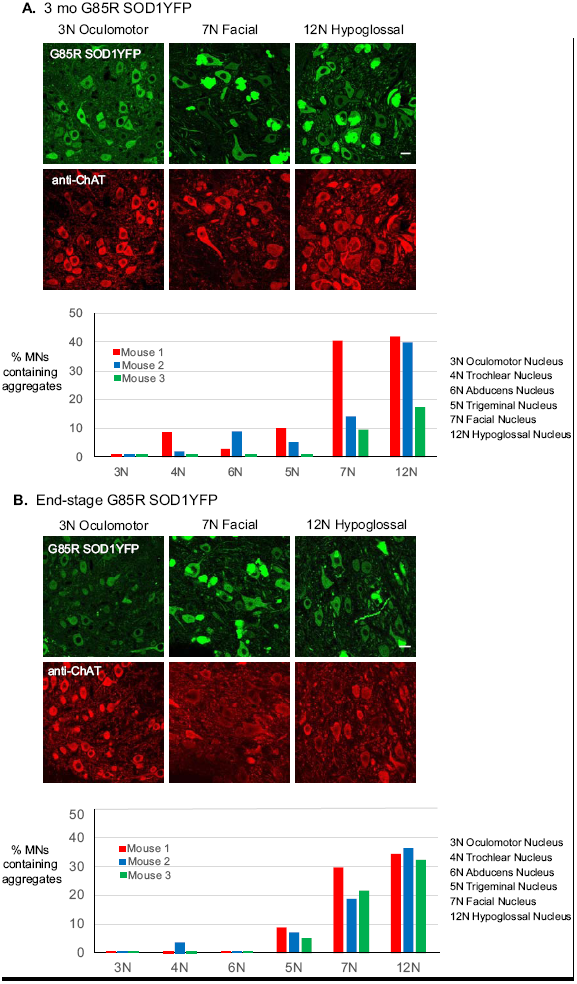
Aggregation of SOD1YFP in cranial nerve motor nuclei. Panels A and B show aggregation in ChAT positive motor neurons from cranial nerve nuclei of G85R SOD1YFP mice at 3 months (panel A) and end-stage (panel B), with representative images shown (top) from 3N (oculomotor), 7N (facial), and 12N (hypoglossal), and bar graphs for 3N, 4N, 6N (extraocular innervation) and 5N, 7N, 12N (other motor nuclei, excluding the motor division of 10N, which did exhibit aggregates). Scale bars are 20 microns and apply to all images. Aggregates are more abundant in the latter group at both 3 months and end-stage, but 5N is less affected and exceeds the extraocular nuclei only at end-stage. Sample preparation, ChAT immunostaining, and microscopy were carried out as in ref. 6. Aggregates were counted in 20 micron coronal sections from 3 mice at 3 months and 3 mice at end-stage. The neuron counts used to generate the bar graphs are shown in Table 1.

**Table 1.** Percentage of motor neurons with aggregates in cranial nerve nuclei of G85R SOD1YFP mice at 3 months of age and at end-stage.

| **3 mo G85R**<br>**SOD1YFP** | **3N** | **4N** | **6N** | **5N** | **7N** | **12N** |
| --- | --- | --- | --- | --- | --- | --- |
| Mouse 1 (cn 298) | 0% (0/79) | 8% (5/59) | 3% (1/36) | 10% (6/60) | 41% (48/118) | 42% (47/112) |
| Mouse 2 (cn 247) | 0% (0/76) | 2% (1/43) | 9% (1/11) | 5% (4/78) | 14% (32/226) | 40% (44/109) |
| Mouse 3 (cn 290) | 0% (0/70) | 0% (0/44) | 0% (0/25) | 0% (0/124) | 10% (24/251) | 17% (17/98) |

| **End stage G85R**<br>**SOD1YFP** | **3N** | **4N** | **6N** | **5N** | **7N** | **12N** |
| --- | --- | --- | --- | --- | --- | --- |
| Mouse 1 (cn 262) | 0% (0/92) | 0% (0/54) | 0% (0/75) | 9% (6/69) | 30% (41/138) | 34% (34/99) |
| Mouse 2 (cn 269) | 0% (0/93) | 4% (2/54) | 2% (1/67) | 7% (6/84) | 19% (27/143) | 37% (37/101) |
| Mouse 3 (cn 268) | 0% (0/90) | 0% (0/22) | 0% (0/62) | 5% (1/22) | 22% (38/175) | 32% (32/99) |

cn = copy number of G85R SOD1YFP transgene.

Could the absence of aggregation in 3N be simply a function of reduced expression of G85R SOD1YFP in motor neurons of that cranial nerve relative to those in sites where aggregation occurs, for example, motor neurons of 12N or spinal cord? We compared levels of mRNA encoding the mutant protein from laser captured motor neurons from 3N, 12N, and spinal cord (SC) of a G85R SOD1YFP mouse (6 wk old, copy number 287) by qRT-PCR and observed that the amount of mRNA was reduced by 15-20% in 3N relative to 12N and spinal cord. Correspondingly, cytosolic YFP fluorescence intensity in 3N motor neurons in perfusion-fixed tissue of two mice with similar copy number was also reduced by 25-30% relative to 12N and spinal cord motor neurons lacking aggregates (see scatter plots of two “low copy” mice in Fig. 2).

**Fig.2.**
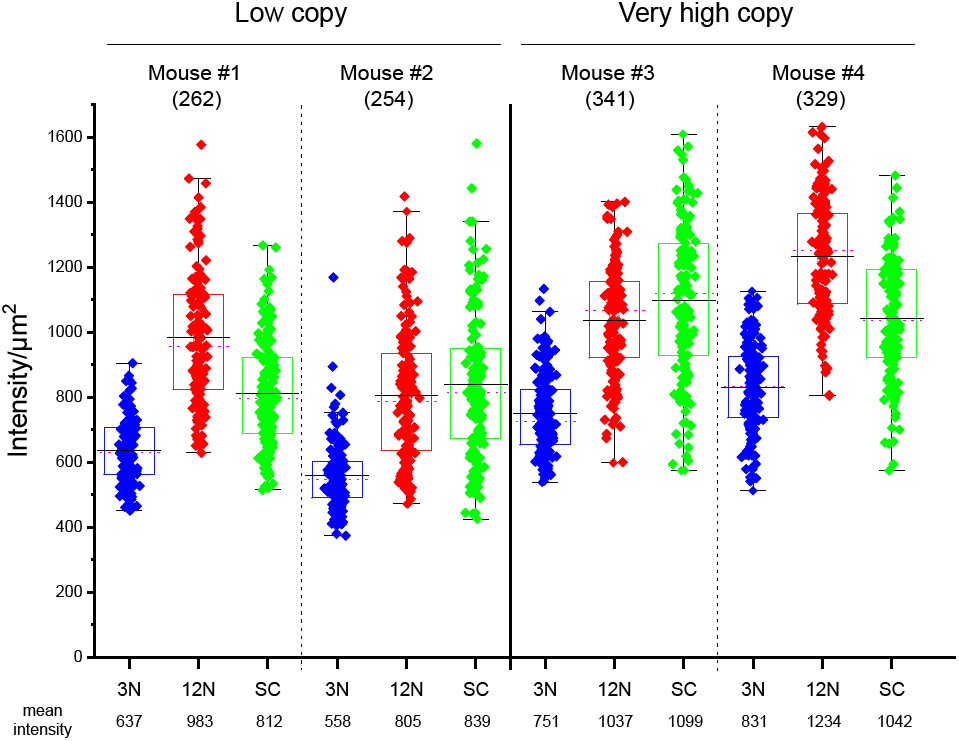
Cytosolic YFP fluorescence intensity of motor neurons from cranial nerves 3N and 12N and from spinal cord from low and very high copy number G85R SOD1YFP transgenic mice. Scatter plots show YFP fluorescence intensities of 150 motor neurons each from oculomotor (3N) and hypoglossal (12N) cranial nerve motor nuclei and spinal cord (SC) of two month old G85R SOD1YFP transgenic mice with “low” and very high copy number, mouse #1,2 and 3,4, respectively. For each tissue, 150 motor neurons (devoid of aggregates in the case of 12N and SC) were analyzed. A 20 μm^2^ region of interest was randomly chosen from 1 μm thick optical (z) sections, and gray scale values were summed for the pixels in that area. The scatter plots encompass all 150 neurons, presented as individual points. The boxes indicate 75^th^ (top) and 25^th^ (bottom) percentiles, the means are indicated at bottom below the tissue designation, and the whiskers (horizontal black lines) indicate 1.5 SD above and below the mean.

Was a 25-30% reduction in the steady-state amount of G85R SOD1YFP, implied by the reduction in fluorescence, in 3N relative to 12N and spinal cord motor neurons sufficient to account for the lack of aggregation in 3N? To address this, we identified two G85R SOD1YFP mice with very high copy numbers, 341 and 329, respectively, exceeding by 25-50% the copy number of animals usually employed in our studies (i.e. ranging 240-280, here termed “low copy”). We observed in these very high copy mice at 2 months of age that, corresponding with the copy number increase, the cytosolic fluorescence of their 3N motor neurons was now increased by ∼20-50% (p<10^−4^; Fig. 2; compare blue plots mouse #s 3&4 with blue plots #s 1&2). The fluorescence intensities in 3N of the very high copy mice nearly completely overlapped within the fluorescence intensities in 12N and SC of two age-matched low copy mice [compare blue scatter plots (3N) of mouse #s 3&4 with red (12N) and green (SC) scatter plots of mouse #s 1&2]. The overlap of the fluorescence intensities was more formally addressed by assignment of a “positional overlapping score” (range 0.0-1.0) (5) to pairs of scatter plots of motor neuron intensities (see Tables 2 and 3). Table 2 shows pairwise comparisons of 3N and 12N sets from within the low copy and high copy mice and between them (sets shown in Fig. 2). Most relevant are the overlap percentage calculations (bold numbers in Table 2) between low copy 3N and low copy 12N (8%, 26%, 27%, 43%) compared with the overlap percentages between high copy 3N and low copy 12N (48%, 55%, 62%, 66%). Clearly, the overlap of sets has been significantly enhanced by increased copy number. A similar increase in the positional overlapping scores are seen when 3N of low copy and high copy are compared with spinal cord low copy (bold numbers in Table 3; 25%, 28%, 43%, 52% vs 56%, 69%, 69%, 81%).

**Table 2.** Proportional overlapping scores comparing motor neuron G85R SODYFP intensities between cranial nerve nuclei in low copy and very high copy mice. Data from Fig. 2 were analyzed by the algorithms in ref. 5. For each pair, the proportional overlapping score is shown, followed by the 95% confidence interval in parentheses. The overlaps highlighted in bold indicate comparisons discussed in the text.

|  | 1-3N | 1-12N | 2-3N | 2-12N | 3-3N | 3-12N | 4-3N | 4-12N |
| --- | --- | --- | --- | --- | --- | --- | --- | --- |
| 1-3N |  | **0.27**<br>(0.20, 0.33) | 0.65<br>(0.58, 0.70) | **0.43**<br>(0.34, 0.57) | 0.68<br>(0.60, 0.73) | 0.12<br>(0.06, 0.18) | 0.47<br>(0.38, 0.53) | 0.01<br>(0.00, 0.02) |
| 1-12N |  |  | **0.08**<br>(0.04, 0.13) | 0.71<br>(0.60, 0.79) | **0.48**<br>(0.37, 0.60) | 0.84<br>(0.68, 0.93) | **0.62**<br>(0.49, 0.71) | 0.61<br>(0.54, 0.67) |
| 2-3N |  |  |  | **0.26**<br>(0.18, 0.36) | 0.37<br>(0.28, 0.43) | 0.02<br>(0.00, 0.05) | 0.19<br>(0.10, 0.24) | 0<br>(0, 0) |
| 2-12N |  |  |  |  | **0.55**<br>(0.44, 0.80) | 0.58<br>(0.44, 0.69) | **0.66**<br>(0.51, 0.85) | 0.32<br>(0.21, 0.42) |
| 3-3N |  |  |  |  |  | 0.33<br>(0.23, 0.47) | 0.76<br>(0.68, 0.85) | 0.10<br>(0.05, 0.19) |
| 3-12N |  |  |  |  |  |  | 0.57<br>(0.45, 0.63) | 0.67<br>(0.60, 0.72) |
| 4-3N |  |  |  |  |  |  |  | 0.24<br>(0.17, 0.31) |
| 4-12N |  |  |  |  |  |  |  |  |

**Table 3.**
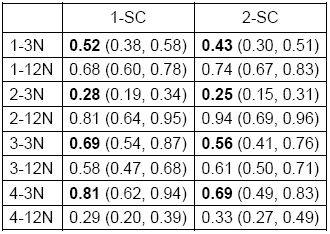
Proportional overlapping scores comparing motor neuron G85R SODYFP intensities between spinal cord and cranial nerve nuclei in low copy and very high copy mice. Data from Fig. 2 were analyzed by the algorithms in ref. 5. For each pair, the proportional overlapping score is shown, followed by the 95% confidence interval in parentheses. The overlaps highlighted in bold indicate comparisons discussed in the text.

Strikingly, despite the florid aggregation in 12N and in SC of the very high copy mice (resembling that shown in Fig. 1), not a single aggregate was found in sections comprising the entirety of 3N from the two very high copy mice. Thus, when 3N expression is increased as in the very high copy mice, reaching a level in most motor neurons that corresponds to that in aggregate-forming 12N and SC motor neurons of low copy mice, there is still no aggregation observed. We thus conclude that 3N motor neuron protection is not a function of the lower level of mutant protein expression in 3N relative to 12N and spinal cord.

We next addressed whether motor neuron survival in cranial nerve nuclei was affected by aggregation, comparing 3N and 12N from 3 mo control ChAT promoter-EGFP mice with G85R SOD1YFP mice at 3 months and end-stage, counting motor neurons in 20 micron coronal sections taken at 100 micron intervals down the length of these nuclei (totaling 5 sections for 3N and 10 sections for 12N; see images in Fig. 3A and 3B for representative sections). The total of ChAT-positive motor neurons was then determined for each of 3 animals from the three groups. As shown in the bar graphs of Fig. 3A, the number of motor neurons in 3N was indistinguishable when comparing ChAT-EGFP with G85R SOD1YFP 3 mo and endstage (the means were 297, 290, 327, non-significant *p* values with t-test). The lack of motor neuron loss in 3N thus mirrors the lack of aggregation. By contrast, the mean motor neuron count for 12N was reduced significantly in end-stage G85R SOD1YFP relative to ChAT-EGFP (Fig. 3B; 507 vs. 708, *p* = 0.01). Thus for 12N, the occurrence of substantial aggregation was associated with significant cell loss, amounting to ∼30%.

**Fig.3.**
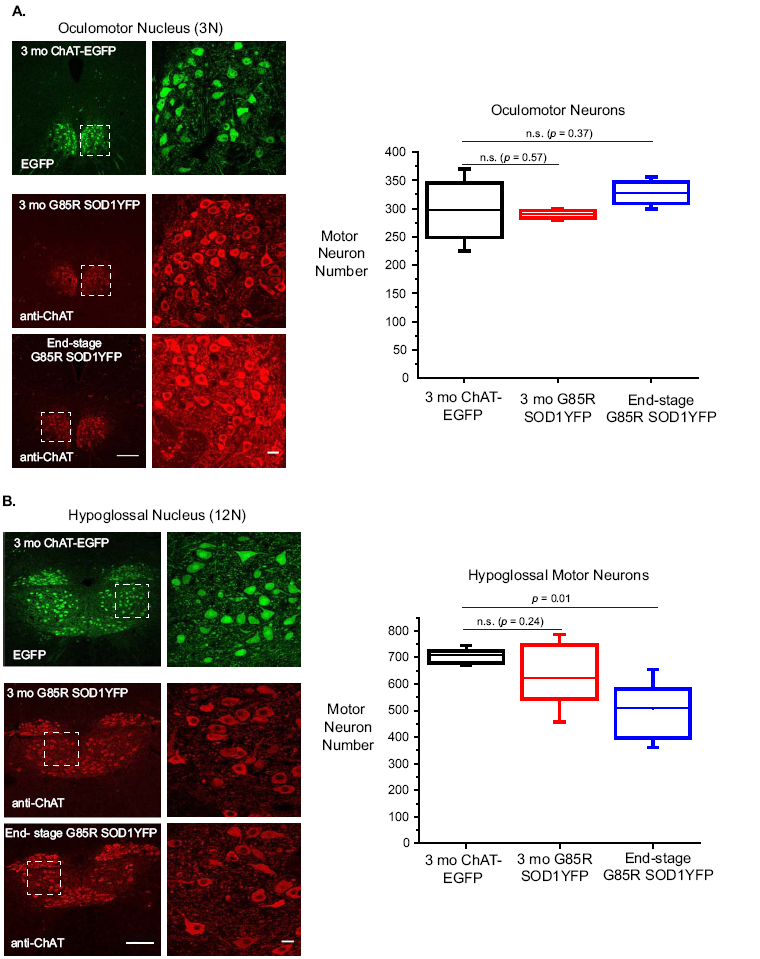
Survival of motor neurons in cranial nerve nuclei 3 and 12 (oculomotor and hypoglossal) of G85R SOD1YFP ALS mice. Panels A and B show ChAT positive motor neurons of 3N and 12N, respectively, determined either from direct GFP fluorescence of ChAT-EGFP mice or anti-ChAT antibody staining of sections from G85R SOD1YFP mice of 3 months or end-stage. At left are low magnification images (scale bar 200 microns) and at right are high resolution images of the indicated areas (scale bar 20 microns). At right are box plots of the motor neuron counts corresponding to the respective mice (n = 3 for each designation). The upper and lower extents of the boxes indicate the 75^th^ and 25^th^ percentiles of the data, respectively. The lines inside the boxes indicate the means, and the whiskers ± 1.5 S.D. *p* values determined by paired t-tests (Origin); n.s., non-significant. ChAT positive motor neurons were counted in 20 micron (coronal) sections taken every 100 microns along the length of the respective nuclei (5 such sections from 3N and 10 from 12N). The sections were stained with anti-ChAT antibody (6), positive cells counted, and counts summed across the sections for the cranial nucleus of a given animal. The numbers were compared with those in the corresponding cranial nuclei of a ChAT promoter-EGFP mouse. 3 mice each of ChAT-EGFP (3 mo), 3 mo G85R SOD1YFP, and end-stage G85R SOD1YFP were analyzed.

In sum, the foregoing data indicate that the cranial nerves supplying extraocular muscles in our SOD1-linked ALS mice are spared of aggregation relative to the other cranial motor nuclei, and where motor neuron survival was scored, there was no cell loss in 3N (oculomotor) vs. significant cell loss in 12N (hypoglossal) by end-stage. Thus, there appears to be a relationship between aggregation and cell loss, albeit it remains to be seen whether the former can drive the latter. It may rather be that the same cellular pathophysiology that drives aggregation can, at the same time, direct cell death. Regardless, the difference in behavior between 3N and 12N can serve as a basis for trying to understand mechanism(s) of protection/sensitivity. We presume that ultimately other mouse models of ALS will likely exhibit the same extraocular motor sparing and that, more generally, at least some of the features of motor collapse in SOD1-linked ALS mice can serve to provide understanding of “sporadic” versions of ALS where, potentially, at least some similar steps of motor neuron loss are occurring.

## Acknowledgments

We thank HHMI for supporting this work. E.V.T. was supported by the Nelson Fund. We are especially grateful to Elizabeth Engle, Children’s Hospital/Harvard Medical School, for suggesting inspection of cranial nuclei in G85R SOD1YFP mice for aggregates.

